# DEBBIES to compare life history strategies across ectotherms

**DOI:** 10.1101/2023.08.22.554265

**Authors:** Isabel M. Smallegange, Sol Lucas

**Author notes:** corresponding author(s): Isabel Smallegange.

## Abstract

Demographic models are used to explore how life history traits structure life history strategies across species. This study presents the DEBBIES dataset that contains estimates of eight life history traits (length at birth, puberty and maximum length, maximum reproduction rate, fraction energy allocated to respiration versus reproduction, von Bertalanffy growth rate, mortality rates) for 185 ectotherm species. The dataset can be used to parameterise dynamic energy budget integral projection models (DEB-IPMs) to calculate key demographic quantities like population growth rate and demographic resilience, but also link to conservation status or biogeographical characteristics. Our technical validation shows a satisfactory agreement between observed and predicted longevity, generation time, age at maturity across all species. Compared to existing datasets, DEBBIES accommodates (i) easy cross-taxonomical comparisons, (ii) many data-deficient species, and (iii) population forecasts to novel conditions because DEB-IPMs include a mechanistic description of the trade-off between growth and reproduction. This dataset has the potential for biologists to unlock general predictions on ectotherm population responses from only a few key life history traits.

## Background & Summary

Matrix population models (MPMs) and integral projection models (IPMs) provide the basis for exploring how variation in the demographic rates of survival, growth and reproduction fuels variation across species in life history traits (like the timing, intensity, frequency and duration of key demographic processes, such as longevity, generation time or degree of iteroparity) and in combinations of life history traits that form life history strategies (including pace of life and reproductive strategy)^1-3^. Life history traits and strategies calculated from these structured population models predict not only key demographic properties such as population growth rate and demographic resilience, but also have important connections to other disciplines like biogeography, evolutionary biology and conservation biology^4-6^.

Important datasets exist that collate MPMs (COMPADRE^7^; COMADRE^8^) and IPMs (PADRINO^9^) for plants and animals, and even algae, fungi, bacteria, and viruses. Recent efforts have furthermore linked the latter datasets in a centralised meta-database of trait data (MOSAIC^10^) so they can be interrogated at the same time. While these datasets are valuable in improving both data access and data usability^11^ and are used globally in networks like the Open Trait Network (https://opentraits.org/), the parameterisation of the structured population models that are within these datasets requires long-term individual-level data that are scored from birth till death. Yet, there are many organisms for which such data are not available, for example, because it is difficult to track individuals over their lifetime (e.g. micro-organisms, small (soil-dwelling) animals). Because such MPM and IPM datasets can form part of pipelines to develop, for example, essential biodiversity variables to observe and report global biodiversity change^12^, it is pertinent to avoid any unwilling species bias. We thus critically need to include datasets of structured population models that can also accommodate more data-deficient species to have a taxonomically most balanced representation as possible.

One type of structured population model that does not require many long-term individual-level life history data is the dynamic energy budget (DEB) IPM^13^. To parameterise a DEB-IPM for a species, one requires eight life history traits to be estimated (traits include length measures, rates of growth, reproduction and mortality [Fig. 1]) to be able to predict survival, growth and reproduction for a simple life cycle (more complex life cycles would require more parameters)^13^. These traits are assumed to be fixed for a life cycle (Fig. 1); that is, length at birth and at puberty represent specific moments in the life cycle and are time-invariant; maximum length and maximum reproduction rate, in turn, are maximum values that are achieved under the most favourable circumstances a life cycle is in, and thus are also time-invariant. We assume that kappa remains constant over a life cycle because empirical evidence suggests so^14^. To estimate the von Bertalanffy growth rate and mortality parameters (Fig. 1) one does require repeated observations on individual growth or survival of individuals within a population. So far, only two small datasets of DEB-IPM parameters have been generated, comprising estimates of the eight life history traits for 13 species of marine megafauna^3^ and 13 microorganisms^15^. Here, we introduce a much larger and taxonomically diverse dataset that we refer to as DEBBIES in which we compiled estimates of the eight life history traits for 185 ectotherm species. Ectotherms are taxonomically diverse and their growth and reproduction can be captured in simple energy budget models, like the one incorporated into the DEB-IPM. Also, more than 99% of species are ectotherms^16^; consequently, no biological prediction can be considered universal if it is does not include these organisms. We find in our technical validation that our model outputs exhibit good agreement with observations on key life history traits (age at maturity, longevity, generation time). The dataset can be used for a variety of different applications of eco-evolutionary studies (e.g.^15,17^) (Fig. 1). Because DEB-IPMs (unlike MPMs and other IPMs) across species are built from the same life history traits, they can readily be used for comparative studies of life histories and population dynamics across a wide range of species, for which DEBBIES provides all the necessary input data. Finally, because DEB-IPMs include a mechanistic description of the trade-off between growth and reproduction, they are particularly suited to create population forecasts to novel conditions (like those created by climate change)^13^ for the species currently listed in the dataset.

**Figure 1.**
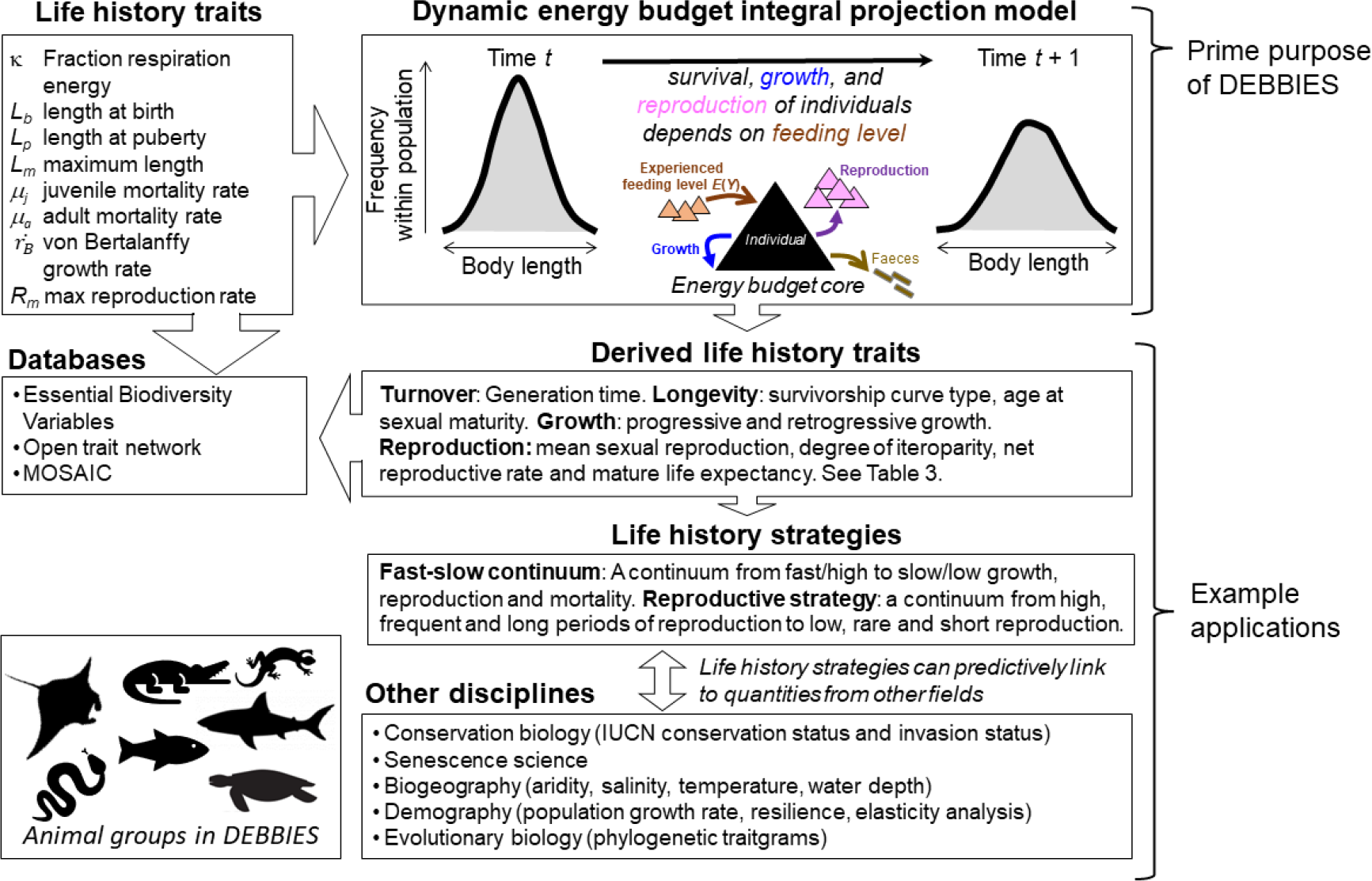
Workflow of parameterising a DEB-IPM and example applications, including what databases DEBBIES can feed into (Essential Biodiversity Variables^12^; MOSAIC^10^). DEBBIES currently contains 185 ectotherms of 18 different orders. Eight life history trait values are required to parameterise a DEB-IPM (top-right box). Once parameterised, it can be used to calculate a further nine derived life history traits (Table 3) that can in turn be summarised into life history strategies^1^. The resulting ‘fast–slow continuum and reproductive strategy framework’ can be linked to quantities from other disciplines^4^.

## Methods

Because the main purpose of DEBBIES is to parameterise DEB-IPMs for further analysis (Fig. 1), we first give a brief description of what a DEB-IPM is, and how a species’ eight life history traits feed into a DEB-IPM, before explaining how we collated the life history data presented in DEBBIES.

### Brief description of a DEB-IPM

The three demographic rates of survival, growth, reproduction and the relationship between parent and offspring size are captured in the DEB-IPM by four fundamental functions, which describe the dynamics of a population comprising cohorts of females of different sizes^3,13^: (1) the survival function, *S*(*L*(*t*)), describing the probability of surviving from time *t* to time *t*+1; (2) the growth function, *G*(*L*^′^, *L*(*t*)), describing the probability that an individual of body length *L* at time *t* grows to length *L’* at *t* + 1, conditional on survival; (3) the reproduction function, *R*(*L*(*t*)), giving the number of offspring produced between time *t* and *t* + 1 by an individual of length *L* at time *t*; and (4) the parent-offspring function, *D*(*L*^′^, *L*(*t*)), the latter which describes the association between the body length of the parent *L* and offspring length *L*’ (i.e. to what extent does offspring size depend on parental size). Denoting the number of females at time *t* by *N*(*L, t*) means that the dynamics of the body length number distribution from time *t* to *t*+1 can be written as:

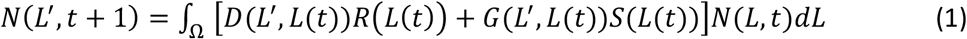

where the closed interval Ω denotes the length domain.

The survival function *S*(*L*(*t*)) in equation (1) is the probability that an individual of length *L* survives from time *t* to *t* + 1:

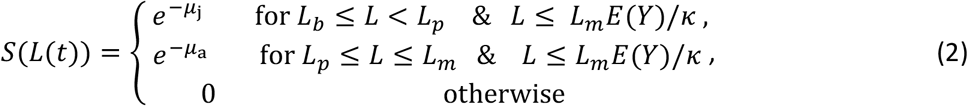

where *E*(*Y*) can range from zero (empty gut) to one (full gut). Individuals die from starvation at a body length at which maintenance requirements exceed the total amount of assimilated energy, which occurs when *L* > *L*_*m*_ · *E*(*Y*)/*κ* and hence, then, *S*(*L*(*t*)) = 0 (e.g., an individual of size *L*_*m*_ will die of starvation if *E*(*Y*) < *κ*). Juveniles and adults often have different mortality rates, and, thus, juveniles (*L*_*b*_ ≤ *L* < *L*_*p*_) that do not die of starvation (i.e. *L* ≤ *L*_*m*_ · *E*(*Y*)/*κ*) have a mortality rate of μ_j_, and adults (*L*_*p*_ ≤ *L* ≤ *L*_*m*_) that do not die of starvation (i.e. *L* ≤ *L*_*m*_ · *E*(*Y*)/*κ*) have a mortality rate of μ_a_.

The demographic functions that describe growth and reproduction in the DEB-IPM are derived from the Kooijman-Metz model^3,13,18^. This is is a simple version of the standard model of Kooijman’s DEB theory, but one that still fulfils the criteria for general explanatory models for the energetics of individuals^19^. The Kooijman-Metz model assumes that individual organisms are isomorphic (which means that body surface area and volume are proportional to squared and cubed length, respectively). The rate at which an individual ingests food, *I*, is assumed to be proportional to the maximum ingestion rate *I*_*max*_, the current feeding level *Y* and body surface area, and hence to the squared length of an organism: *I* = *I*_*max*_*YL*^2^. Ingested food is assimilated with a constant efficiency *ε*. A constant fraction *κ* of assimilated energy is allocated to respiration; this respiration energy equals *kεI*_*max*_*YL*^2^ and is used to first cover maintenance costs, which are proportional to body volume following *ξL*^3^ (*ξ* is the proportionality constant relating maintenance energy requirements to cubed length), while the remainder is allocated to somatic growth. The remaining fraction 1 – *κ* of assimilated energy, the reproduction energy, is allocated to reproduction in case of adults and to the development of reproductive organs in case of juveniles, and equals (1 − *κ*)*εI*_*max*_*YL*^2^. This means that, if an individual survives from time *t* to time *t*+1, it grows from length *L* to length *L*’ following a von Bertalanffy growth curve, 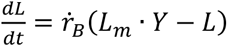, where 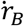 is the von Bertalanffy growth rate (here assumed to be constant across feeding levels, but can be adjusted if necessary^13^) and *L*_*m*_ = *kεI*_*max*_/*ξ* is the maximum length under conditions of unlimited resource. Both *κ* and *I*_*max*_ are assumed to be constant across experienced feeding levels, and therefore *L*_*m*_ is also assumed constant.

Implicitly underlying the population-level model of equation (1), like in any IPM, is a stochastic, individual-based model, in which individuals follow Markovian growth trajectories that depend on an individual’s current state^20^. This individual variability is in standard IPMs modelled in the functions describing growth, *G*(*L*^′^, *L*(*t*)), and the parent-offspring association, *D*(*L*^′^, *L*(*t*)) (see below), using a probability density distribution, typically Gaussian^21^. In the DEB-IPM, this individual variability arises from how individuals experience the environment; specifically, the experienced feeding level *Y* follows a Gaussian distribution with mean *E*(*Y*) and standard deviation *σ*(*Y*). It means that individuals within a cohort of length *L* do not necessarily experience the same feeding level due to demographic stochasticity (e.g. individuals, independently of each other, have good or bad luck in their feeding experience). Taken together, this means that the function *G*(*L*^′^, *L*(*t*)) is the probability that an individual of body length *L* at time *t* grows to length *L’* at *t* + 1, conditional on survival, and, following common practice^20-22^, follows a Gaussian distribution:

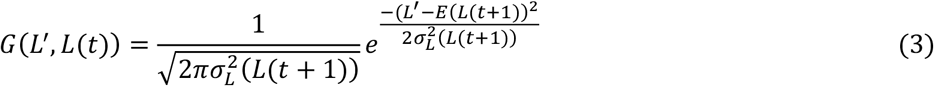

with the growth realized by a cohort of individuals with length *L*(*t*) equalling

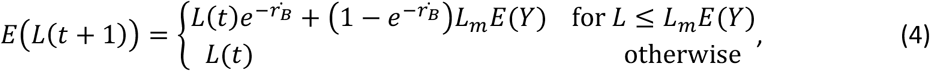

and the variance in length at time *t* + 1 for a cohort of individuals of length *L* as

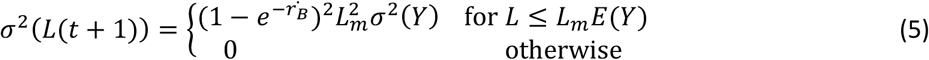

where *σ*(*Y*) is the standard deviation of the expected feeding level.

If a surviving female is an adult, she also produces offspring. According to the Kooijman-Metz model^18^, reproduction, i.e. the number of offspring produced by an individual of length *L* between time *t* and *t* + 1, equals 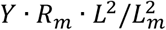 . The parameter *R*_*m*_ is the maximum reproduction rate of an individual of maximum length *L*_*m*_. Note that *R*_*m*_ is proportional to (1 – *κ*)^18^, whereas *L*_*m*_ is proportional to *κ*; *κ* thus controls the trade-off between energy allocation to reproduction versus growth. However, the role of *κ* in the DEB-IPM is mostly implicit, as *κ* is used as input parameter only in the starvation condition (see below), whereas *R*_*m*_ and *L*_*m*_ are measured directly from data. Like *L*_*m*_, *R*_*m*_ is also proportional to *I*_*max*_; since both *κ* and *I*_*max*_ are assumed to be constant across experienced feeding levels, *R*_*m*_ is also assumed constant. The reproduction function *R*(*L*(*t*)) gives the number of offspring produced between time *t* and *t* + 1 by an individual of length *L* at time *t*:

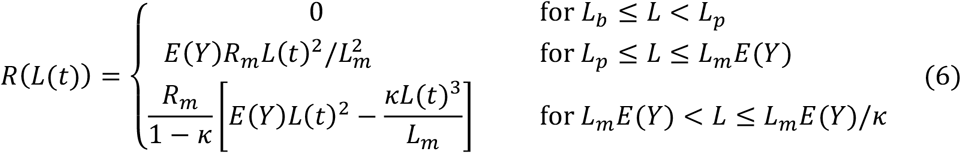

Individuals are mature when they reach puberty at body length *L*_*p*_ and only surviving adults reproduce (equation (1)); thus, only individuals within a cohort of length *L*_*p*_ ≤ *L* ≤ *L*_*m*_*Y*/*k* reproduce.

The probability density function *D*(*L*^′^, *L*(*t*)) gives the probability that the offspring of an individual of body length *L* are of length *L’* at time *t* + 1, and hence describes the association between parent and offspring character values:

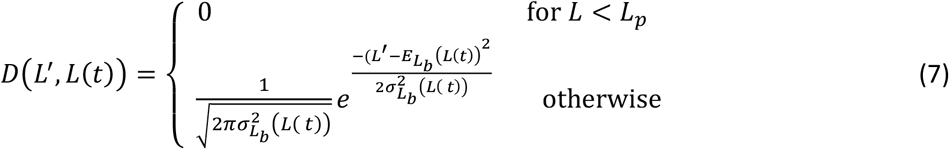

where 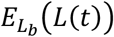 is the expected size of offspring produced by a cohort of individuals with length *L*(*t*), and 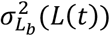 the associated variance. For simplicity, we set 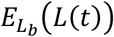 constant and assumed its associated variance, 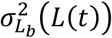, to be very small.

Finally, note that the DEB-IPM assumes no effect of temperature on fundamental functions. Temperature effects on fundamental functions, however, can be approximated by varying experienced feeding level. Alternatively, one could resort to a more detailed and more parameter-rich DEB-IPM that links individuals’ size- and temperature-dependent consumption and maintenance via somatic growth, reproduction, and size-dependent energy allocation to emergent population responses^23^, but this is not linked to DEBBIES.

### Collection of life history trait data required to parameterise a DEB-IPM

Running a DEB-IPM for a species requires estimates for eight life history traits: the fraction respiration energy *κ*, length at birth *L*_*b*_, length at puberty *L*_*p*_, maximum length *L*_*m*_, juvenile mortality rate μ_j_, adult mortality rate μ_a_, von Bertalanffy growth rate 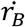 and maximum reproduction rate *R*_*m*_ (Fig. 1).

For the elasmobranchs, we obtained von Bertalanffy growth rate values using the following search, in order of priority: (i) primary literature, using female growth curve, measured empirically using data from animals, (ii) from Froese^24^ supplementary material, or (iii) using the equation 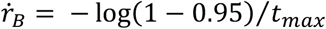, where *t*_max_ is a species’ longevity (years), sourced from Fishbase^25^. If multiple values were available, only those in the highest priority group were kept. If there were still multiple values, their median value was used. Any values listed as ‘Questionable’ on Fishbase^25^ were not used. Body lengths for 152 species were sourced from Sharks of the World^26^ or Rays of the World^27^. Total lengths were converted to fork lengths using scalar values on Fishbase^26^. Length at birth was sourced from the IUCN red list^28^ for five species (*M. ambigua, M. birostris, A. parmifera, D. trachyderma* and *R. australiae*). Length at puberty for *Maculabatis ambigua* was sourced from primary literature^29^. We calculated maximum reproduction rate *R*_*m*_ as *R*_*m*_ = (*c* × *n*)/*i*, where *c* is the mean clutch size, *n* is the mean number of litters produced per year, and *i* is the remigration interval, which is the minimal number of years between reproductive seasons.

Minimum and maximum pup numbers were sourced from Sharks of the World^26^, Rays of the World^27^, IUCN red list^28^, Fishbase^25^, or Barrowclift et al.^30^. The sources for maximum and minimum pup numbers were not prioritised, and therefore some maximums and minimums were obtained from different sources. Breeding intervals were found in the same way, except for *C. granulosus, S. californica, F. macki* and *R. porosus*, which data we sourced from primary literature^31-34^. If the number of pups or breeding interval were not found, we took those data from the next closest species within the same genus. Mode of reproduction was sourced from the Sharks of the World^26^ and Rays of the World^27^. Adult mortality rate was calculated^35^ by taking the inverse of the mean of longevity and age at maturity (*a*) (*μ*_*a*_ = 1/[(*t*_*max*_ + *a*)/^2^]. Juvenile mortality rate^36^ was calculated as 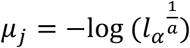, where survival to maturity^37^, *l*_*α*_, equals 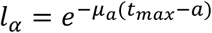.

Values for all other species were taken directly from the primary literature. Finally, for most species, we were able to take values for *κ* from the Add My Pet database^38^. If no values for *κ* were available for a species, we assumed *κ* = 0.8 as is common practice^39^. The parameter kappa is explicit in the starvation condition^13^, which states that individuals of length *L* die of starvation when they can no longer cover their maintenance costs, which occurs when *L* = *L*_*m*_*E*(*Y*)*κ* = *L*_starvation_. At any feeding level, the ultimate length that individuals can grow to equals *L*_∞_ = *L*_*m*_*E*(*Y*). Substituting *L*_∞_ into the starvation condition returns: *L*_starvation_ = *L*_∞_/*κ*. In our case, *κ* = 0.8 so that *L*_starvation_ = 1.^2^5*L*_∞_. This means that at any constant feeding level, individuals will never reach the length at which they starve because that is 25% larger than the ultimate length they can achieve at that feeding level. Only when feeding level varies over time, like in a stochastic time series, can *κ* affect population responses if, for example, individuals that were large in a good feeding environment suddenly find themselves in a poor feeding environment.

## Data Records

The DEBBIES dataset (Version 5) includes eight life history trait records for 185 ectotherm species that were sourced from the scientific literature and are stored in one csv file with accompanying metadata text file on FigShare^40^. Each row in the file represents one species record. For most species, we sourced life history traits from different studies as rarely only one single study provided estimates for all traits for one species. Each data record therefore describes a species’ general life history and is not specific to a particular population. On the one hand, this approach is in line with the assumption of DEB theory that individual-level differences are sufficiently small to take mean values to represent species-specific parameters^41^. On the other hand, recent work on different populations of Trinidadian guppies cautions against combining life history data from different studies, because systematic bias during parameter estimation can generate substantial variation, and similar patterns of growth and reproduction can be produced with very different parameter sets^42^. In contrast, focusing only on parameter sets specific to a particular population of Trinidadian guppies returns life history variation predictions that are in line with observations^42^. Users should thus carefully interpret their results, especially when predictions are not in line with expectations. One could, for example, explore the robustness of a DEB-IPM to uncertainty or perturbation of the input data to assess how much the data can be changed before any desired property of the model, like predicted population growth rate, is lost^43,44^. The columns of the data file are described in Table 1.

**Table 1.** Description of the columns in the data records file; each row in the file is a species.

| Name | Column | Description |
| --- | --- | --- |
| RecordID | 1 | Record ID |
| Class | 2 | Species class |
| Order | 3 | Species order |
| Family | 4 | Species family |
| Species | 5 | Species Latin name |
| Common_name | 6 | Species common name |
| $\kappa$ | 7 | Fraction of energy allocated to respiration ($\kappa$) (as opposed to reproduction) |
| $L_b$ | 8 | Length at birth (cm) |
| $L_p$ | 9 | Length at puberty (cm) |
| $L_m$ | 10 | Maximum length (cm) |
| $\mu_j$ | 11 | Juvenile mortality rate ($y^{-1}$) |
| $\mu_a$ | 12 | Adult mortality rate ($y^{-1}$) |
| $r_B$ | 13 | von Bertalanffy growth rate ($y^{-1}$) |
| $R_m$ | 14 | Maximum reproduction rate ($y^{-1}$) |
| Contributor | 15 | Name of the person who collected the trait data |
| kappa_REF | 16 | Reference for $\kappa$ |
| $L_b$_REF | 17 | Reference for $L_b$ |
| $L_p$_REF | 18 | Reference for $L_p$ |
| $L_m$_REF | 19 | Reference for $L_m$ |
| $\mu_j$_REF | 20 | Reference for $\mu_j$ |
| $\mu_a$_REF | 21 | Reference for $\mu_a$ |
| $r_B$_REF | 22 | Reference for $r_B$ |
| $R_m$_REF | 23 | Reference for $R_m$ |

## Technical Validation

Two model performance tests on a subset of DEBBIES that have validated the reliability of the dataset are presented elsewhere^3,13^. Here, we conducted a similar model performance test on the full dataset by exploring the distribution of predicted population growth rates, and by comparing predicted and observed generation time (years), age at maturity (years), and longevity (the sum of age at maturity and mature life expectancy; years) for three different feeding levels. Because not all species were represented at all feeding levels (less than half of all species in the dataset can persist at low feeding levels), any effect of feeding level cannot be estimated independently. Therefore, we ran the model performance tests separately for each feeding level. Predicted values of the latter quantities are calculated as explained below in the usage notes (see also Fig. 1: derived life history traits). For ray-finned fish, observed values were obtained from Fishbase^25^ and, if unavailable, we obtained generation times from the IUCN Red List^28^ and age at maturity values from the Animal Diversity Web^45^. For the cartilaginous fish, we obtained generation times (for all species) and longevities (18 species) from the IUCN Red List^28^, and longevity and age at maturity values from the Sharks of the World book^26^, the Rays of the World book^27^ or from Barrowclift et al.^30^. For one cartilaginous fish, we obtained the longevity value directly from the scientific literature (silvertip shark *Carcharhinus albimarginatus*)^47^ and for the Galapagos shark *Carcharhinus galapagensis* from Fishbase^25^. For all other species, we first consulted the Animal Diversity Web^45^, then the IUCN Red List^28^ for observations on generation time and AnAge^47^ for observations on age at maturity and/or longevity if these were unavailable in the Animal Diversity Web^45^. If, for a particular quantity, a value range was given, we took the median; if a series of observations was given, we took the mean.

Our validation shows that the predicted population growth rate *λ* (calculated as the dominant eigenvalue of the matrix approximation of equation (1), see also Table 3) most species are slightly higher than *λ* = 1 (denoting population increase) at high experienced feeding level (*E*(*Y*) = 0.9), centred around *λ* = 1 (denoting stability) at an intermediate feeding level (*E*(*Y*) = 0.7), and mostly lower than *λ* = 1 (denoting population decline) at low feeding level (*E*(*Y*) = 0.5) (Fig. 2a). This is in line with the general expectation in ecology that populations increase under favourable conditions but decline when conditions deteriorate. Predicted generation times were higher than observed generation times at the lower experienced feeding levels but not significantly different from observed generation times at the highest experienced feeding level (Table 2; Fig. 2b). Using the root mean square error (RMSE) (Table 2), we can quantify the overall deviation. Specifically, we estimate that 95% of the observed generation time values fall within a range that extends 19 – 23 years from the predicted generation times across feeding levels (assuming the residuals follow a Normal distribution, 95% of observed values fall within ±2 × RMSE from the predicted values, and we took the lowest and highest RMSE value from across the range of feeding levels to estimate this 95% confidence interval). The average RMSE value for generation time across the three feeding levels equals 10 years (Table 2). Given the fact that the highest observed generation time is 53 years (*Carcharodon carcharias*), the average RMSE of 10 years indicates that the model predictions have an average error rate of 19% (10 ÷ 53 = 0.19). We surmise that one reason for this relatively large error rate is that predicted generation time is, following convention [1], calculated as *T* = log(*R*_0_) /log(*λ*), where *R*_0_ is the net reproductive rate and *λ* the population growth rate. Calculated this way, generation time represents the time it takes a population to increase by a factor *R*_0_, which might not always be a good approximation of how generation time is measured in the field.

**Table 2.** Technical validation that fitted linear regression models without an intercept (*y* ∼ *x*) on observed (*x*) and predicted (*y*) values of generation time (*G*), longevity (*L*) and age at maturity (*A*) for three experienced feeding levels *E*(*Y*) = 0.5, *E*(*Y*) = 0.7, and *E*(*Y*) = 0.9. The 95% confidence intervals (CIs) of the regression coefficient are approximated as the regression coefficient ± twice its standard error. If a 95% CI overlaps with 1, predicted values do not significantly differ from observed values. Also given is the root mean square error (RMSE) and the coefficient of determination (R_2_) of each model. See also Fig. 2.

| $E(Y)$ | model $y \sim x$ | regression coefficient | 95% CI | RMSE | $R^2$ |
| --- | --- | --- | --- | --- | --- |
| 0.5 | $G_{\text{predicted}} \sim G_{\text{observed}}$ | 1.24 | 1.10 – 1.38 | 9.34 | 0.27 |
| 0.7 | $G_{\text{predicted}} \sim G_{\text{observed}}$ | 1.23 | 1.16 – 1.30 | 11.40 | 0.28 |
| 0.9 | $G_{\text{predicted}} \sim G_{\text{observed}}$ | 1.02 | 0.97 – 1.07 | 9.92 | 0.38 |
| 0.5 | $L_{\text{predicted}} \sim L_{\text{observed}}$ | 0.59 | 0.57 – 0.61 | 6.14 | 0.55 |
| 0.7 | $L_{\text{predicted}} \sim L_{\text{observed}}$ | 0.84 | 0.81 – 0.87 | 11.20 | 0.40 |
| 0.9 | $L_{\text{predicted}} \sim L_{\text{observed}}$ | 0.99 | 0.93 – 1.05 | 21.50 | 0.16 |
| 0.5 | $A_{\text{predicted}} \sim A_{\text{observed}}$ | 1.25 | 1.21 – 1.29 | 5.70 | 0.49 |
| 0.7 | $A_{\text{predicted}} \sim A_{\text{observed}}$ | 1.30 | 1.26 – 1.34 | 5.51 | 0.57 |
| 0.9 | $A_{\text{predicted}} \sim A_{\text{observed}}$ | 1.45 | 1.40 – 1.50 | 6.60 | 0.57 |

**Table 3.** Nine key derived life history traits that inform on a species’ turnover rate, longevity, growth and reproduction, for which we provide MatLab code to calculate them (see Code Availability)^40^. To calculate history traits, one needs to discretise the IPM (equation (1)) by dividing the length domain Ω into very small width discrete bins (we chose 200 bins), resulting in a matrix **A** of size *p* × *q*, where *p* = *q* = 200, and which dominant eigenvalue equals λ, the population growth rate. Mean lifetime reproductive success *R*_0_ is the dominant eigenvalue of the matrix **F** = **V**(**I** − **GS**)^−1^, where **I** is the identity matrix and **V** = **DR**, With **D** as the parent-offspring association, **R** the reproduction, **G** the growth and **S** the survival matrix^52^; the gives generation time^52^ *T* = log(*R*_0_)/log(*λ*). The mean life expectancy, *η*_e_, is calculated as *η*_e_ = **1**^T^**N**, where **1** is a vector of ones of length *m* and **N** is the fundamental matrix **N** = (**I** − **S**)^−1^. The longevity of an individual of length *L* is *η*_L_, which means we can calculate age at sexual maturity 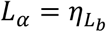and mature life expectancy 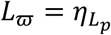 so that 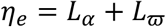 ^48^. *l*_x_ is the probability of surviving to age at least *x*, and *m*_*x*_ is the average fertility of age class *x* (cf.^53^). 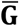 is the mean of **G** 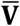 is the mean of **V**, and *i* and *j* are the row and column entries of the matrix, respectively. The vital rates included in progressive growth *γ*, retrogressive growth *ρ*, and sexual reproduction *φ*, were averaged across the columns *j* (the length bins), weighted by the relative contributions of each stage at stationary equilibrium. For example, to calculate mean sexual reproduction *φ*, we summed the values in the columns *j* of the **V** matrix and multiplied each *φ*_*ij*_ by the corresponding *j*th element *w*_j_ of the stable stage distribution **w**, calculated as the right eigenvector of **A**. Finally, the degree of iteroparity 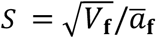, with 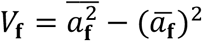, where 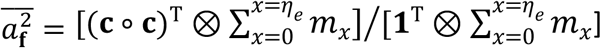 and 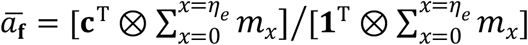, with **c**^T^(1 ^2^ ··· *p*) ^48^.

| Life history trait | Symbol | Definition | Equation |
| --- | --- | --- | --- |
| Generation time | $T$ | Number of days required for the individuals of a population to be fully replaced by new ones | $T = \frac{\log(R_0)}{\log(\lambda)}$ |
| Survivorship curve | $H$ | Keyfitz' entropy ($H < 1$ denotes increasing mortality rate with age, $H > 1$ denotes decreasing mortality rate with age, $H = 1$ denotes a constant mortality rate with age)<sup>54</sup>. | $H = - \frac{\sum_{x=0}^{x=\eta_e} \log(l_x) l_x}{\sum_{x=0}^{x=\eta_e} l_x}$ |
| Age at maturity | $L_\alpha$ | Number of days that it takes an average individual in the population to become reproductive | $L_\alpha = \eta_{L_b}$ |
| Progressive growth | $\gamma$ | Mean probability of growing to a larger length across the length domain $\Omega$. | $\gamma = \sum_i^m \bar{\mathbf{G}}_{i,j} \vert _{i < j}$ |
| Retrogressive growth | $\rho$ | Mean probability of growing to a smaller length across the length domain $\Omega$. | $\rho = \sum_i^m \bar{\mathbf{G}}_{i,j} \vert _{i > j}$ |
| Mean recruitment success | $\varphi$ | Mean per-capita number of recruits across the length domain $\Omega$. | $\varphi = \sum_i^m \bar{\mathbf{V}}_{i,j}$ |
| Degree of iteroparity | $S$ | Coefficient of variation in age at reproduction | $S = \sqrt{V_f}/\bar{a}_f$ |
| Net reproductive rate | $R_0$ | Mean number of recruits produced during the mean life expectancy of an individual in the population. | $R_0 = \sum_{x=0}^{x=\eta_e} l_x m_x$ |
| Mature life expectancy | $L_\omega$ | Number of days from the mean age at maturity ($L_\alpha$) until the mean life expectancy ($\eta_e$) of an individual in the population. | $L_\omega = \eta_{L_p}$ |

**Figure 2.**
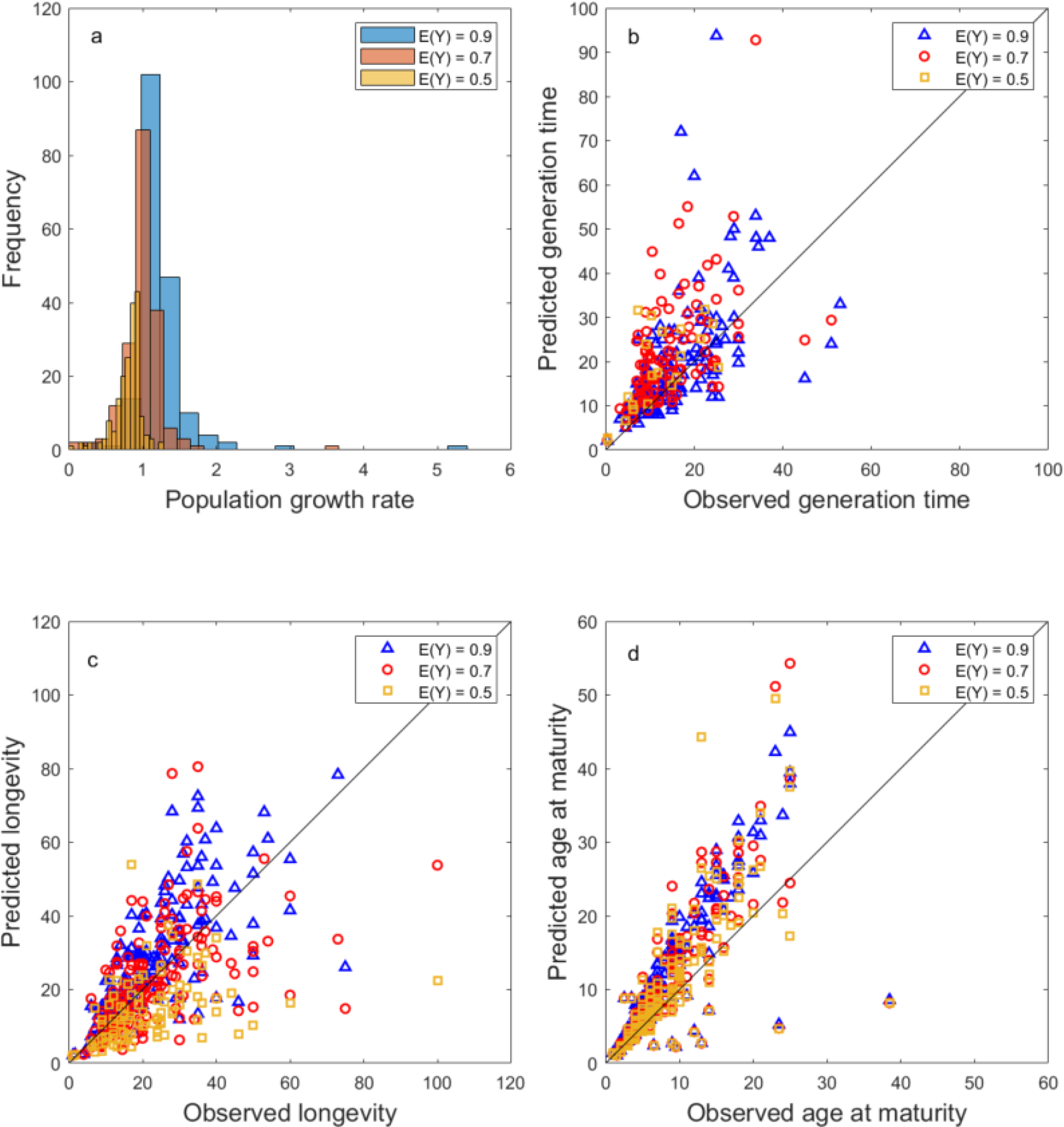
Technical validation. Shown are for three levels of experienced feeding level (blue: *E*(*Y*) = 0.9; red: *E*(*Y*) = 0.7; yellow: *E*(*Y*) = 0.5) the frequency distributions of the population growth rate *λ* (**a**), observed versus predicted generation time (years) (**b**); observed versus predicted longevity (years) (**c**); and observed versus predicted age at maturity (years) (**d**). The black lines in (B-D) denote the *x* = *y* line of equal values.

Predicted longevity was lower than observed longevity at the lower experienced feeding levels but not significantly different from observed values at the highest experienced feeding level (Table 2; Fig. 2c). Using the root mean square error (RMSE) (Table 2), we estimate that 95% of the observed longevity values fall within a range that extends 12 – 43 years from the predicted longevities across feeding levels. The average RMSE value for longevity across the three feeding levels equals 13 years (Table 2). Given the fact that the highest observed longevity is 100 years (*Squalus suckleyi*), the average RMSE of 13 years indicates that the model predictions have an average error rate of 13% (13 ÷ 100 = 0.13).

Finally, predicted age at maturity was higher than observed age at maturity and the best fit with observed values was at the lowest experienced feeding level (Table 2; Fig. 2d). We are unsure why this is the case, but it most likely indicates a mismatch between the functional biology of maturation, and the assumptions underlying the calculations of age at maturity^48^. Using the root mean square error (RMSE) (Table 2), we estimate that 95% of the observed age at maturity values fall within a range that extends 11 – 13 years from the predicted ages at maturity across feeding levels. The average RMSE value for longevity across the three feeding levels equals 6 years (Table 2). Given the fact that the highest observed longevity is 39 years (*Chelonia mydas*), the average RMSE of 13 years indicates that the model predictions have an average error rate of 15% (6 ÷ 39 = 0.15).

In summary, the model predicts generation times and longevity values accurately at high experienced feeding levels, but predicted generation times showed the highest average error rate. Predicted age at maturity values were significantly higher than observed values. However, their error rate was overall lower than the error rate of predicted generation times and only slightly higher than the error rate of predicted longevity values. Depending on the specific question a user is interested in, these error rates can be acceptable or not. What gives us confidence in the technical quality of the dataset and its potential applications, is the fact that predicted population growth rates are within the range that we would expect biologically (Fig. 1a).

## Usage Notes

This data descriptor was peer reviewed in 2023 based on version 5 of the DEBBIES dataset^40^. All versions are available online^40^ as described as in Table 1. Version 1 contains data on 47 species; some of these entries were incorrect and removed in version 2. In version 3, 157 species of elasmobranchs were added, with three more elasmobranchs added in version 4. In the current version 5, we changed the symbols in the dataset that denote *κ* and 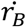 to match the symbols used in this data descriptor. We anticipate future versions to include more species.

The main purpose of the DEBBIES dataset is to be able to parameterise a DEB-IPM for each of the species in the dataset, but the dataset on its own can also feed into other databases (Fig. 1). Once parameterised, running a DEB-IPM requires setting a value for the experienced feeding level *E*(*Y*), which value can range from zero (empty gut) to one (full gut). We find that for many species, *E*(*Y*) should be higher than about 0.7 for the model to run, although it can run for some species at much lower values, down to 0.4^13^. A feeding level of around 0.7 can be considered to represent a gut that is ‘just filled’, on a scale between empty and bursting^49^. In that sense, we should perhaps not be surprised DEB-IPMs require a minimum experienced feeding level of around 0.7.

Many different quantities can be calculated from a parameterised DEB-IPM. For example, like current MPM and IPM datasets^7-9^, DEB-IPMs can be used to calculate key demographic quantities such as population growth rate and demographic resilience, but also nine key derived life history traits that inform on a species’ turnover rate, longevity, growth and reproduction (Fig. 1) (Table 3)^1^. Additionally, because all DEB-IPMs have the same structure, one can run perturbation analyses to estimate for each species the proportional change in the population growth rate for a proportional change in each of the input life history traits (*κ, L*_*b*_, *L*_*p*_, *L*_*m*_, μ_j_, μ_a_, 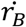, *R*_*m*_). These results can be used to pinpoint those parts of an organism’s life history that should be the focus of monitoring or research to inform conservation management, or those life history traits that contribute most to fitness. We provide MatLab code to calculate the aforementioned quantities (see Code Availability below). Some users, however, might be more interested in exploring the dynamics of single species, rather than performing cross-taxonomical analyses. For those, we have included code to explore how the aforementioned quantities vary with experienced feeding level *E*(*Y*) for a particular species (see Code Availability below).

The nine derived life history traits^1^ (Fig. 1, Table 3) can be used for more in-depth cross-taxonomical analyses (Fig. 1). For example, statistical analyses (e.g., a phylogenetically corrected principal component analysis^50^ can be used to explore how life history variation can be structured and summarised into a few life history strategies^51^. Previous studies using MPM databases have revealed that plant and animal life history variation is predominantly structured along a fast-slow life history speed axis (rapid growth, high reproduction and short lifespan versus slow growth, low reproduction but long lifespan) and a reproductive strategy axis (concentrated versus more dispersed reproduction events across the adult lifespan)^1-2^. An interesting exercise would be to see if the same structuring holds when individual growth and reproduction rates are described mechanistically using an energy budget model (like in DEB-IPMs) instead of being estimated from observational data (like in MPMs). What is more, because a DEB-IPM is run for a specific feeding level set by the user, one could also investigate if the same structuring applies across a range of feeding levels (but note that our validation only covers feeding levels between 0.5-0.9). Finally, if species can be ranked along one or two life history strategy axes, their position along these axes can be linked to quantities from other disciplines (Fig. 1). This opens op possibilities for users to explore the potential for life history strategies to, for example, predict the likelihood of extinction or invasion (e.g.^4^) investigate to what extent such strategies map onto phylogenetic trees (traitgrams) to answer questions about their evolution (e.g.^6^).

Finally, users should be aware that, for the species currently included in DEBBIES, the projection interval between time *t* and time *t* + 1 is set at 1 year; all rates are therefore expressed per year and all durations are expressed in years. Likewise, all length measurements in DEBBIES are currently expressed in centimetres. Future versions, however, could include species which demographic rates are better projected over shorter time intervals and which lengths are better expressed in millimetres or even micrometres (e.g., microorganisms^6^).

## Code Availability

MatLab code and instructions in a README text file for cross-taxonomical analysis can be downloaded from FigShare (https://doi.org/10.6084/m9.figshare.13241972; folder: ‘Derived life history traits across ectotherms’^40^) to calculate (i) generation time, survivorship curve, age at sexual maturity, progressive growth, retrogressive growth, mean sexual reproduction, degree of iteroparity, net reproductive rate, and mature life expectancy (Table 3), as well as population growth rate and demographic resilience (damping ratio), and (ii) run an elasticity analysis for each species listed in DEBBIES for a predefined, experienced feeding level.

For users interested in exploring the dynamics of single species, MatLab code and instructions in a README text file can be downloaded from FigShare (https://doi.org/10.6084/m9.figshare.13241972; folder: ‘Derived life history traits single ecotherm’^40^) to calculate (i) generation time, survivorship curve, age at sexual maturity, progressive growth, retrogressive growth, mean sexual reproduction, degree of iteroparity, net reproductive rate, and mature life expectancy (Table 3), as well as population growth rate and demographic resilience (damping ratio), and (ii) run an elasticity analysis for a single species for a range of experienced feeding levels defined by the user.

## Acknowledgements

The authors would like to acknowledge Gemma Crawford, Naomi Eeltink, Tom Hopman and Iris van Rijn for their help in collecting some of the data as part of their studies, and thank Amy Attle and Burkeleigh Boyd for testing the code as part of their studies. For the purpose of Open Access, the author has applied a CC BY public copyright licence to any Author Accepted Manuscript (AAM) version arising from this submission.

## Author contributions

Isabel Smallegange - project conception – data acquisition – data validation – writing. Sol Lucas - data acquisition – data validation – proof reading.

## References

1. Salguero-Gómez, R., et al. Fast–slow continuum and reproductive strategies structure plant life-history variation worldwide. Proc. Natl Acad. Sci. U.S.A. 113, 230–235 (2016).

2. Paniw, M., Ozgul, A., & Salguero-Gómez, R. Interactive life-history traits predict sensitivity of plants and animals to temporal autocorrelation. Ecol. Lett. 21, 275–286 (2018).

3. Smallegange, I. M., Flotats Avilés, M., Eustache, K. Unusually paced life history strategies of marine megafauna drive atypical sensitivities to environmental variability. Front. Mar. Sci. 7, 597492 (2020).

4. Salguero-Gómez, R. Applications of the fast–slow continuum and reproductive strategy framework of plant life histories. New Phytol. 213, 1618–1624 (2017).

5. Capdevila, P., et al. Longevity, body dimension and reproductive mode drive differences in aquatic versus terrestrial life history strategies. Funct. Ecol. 34, 1613–1625 (2020).

6. Romeijn, J., Smallegange, I. M. Exploring how the fast-slow pace of life continuum and reproductive strategies structure microorganism life history variation. Preprint at 10.1101/2022.11.28.517963 (2022).

7. Salguero-Gómez, R., et al. The COMPADRE Plant Matrix Database: an online repository for plant population dynamics. J. Ecol. 103, 202–218 (2014).

8. Salguero-Gómez, R., et al. COMADRE: a global database of animal demography. J. Anim. Ecol. 85, 371–384 (2016).

9. Levin, S., et al. PADRINO. Zenodo v0.0.1. 10.5281/zenodo.6573870 (2022).

10. Bernard, C., et al. MOSAIC: a unified trait database to complement structured population models. Sci. Data 10, 335 (2023).

11. Gallagher, R. V., et al. Open Science principles for accelerating trait-based science across the Tree of Life. Nature Ecol. Evol. 4, 294–303 (2020).

12. Kissling, W. D., et al. Towards global data products of Essential Biodiversity Variables on species traits. Nature Ecol. Evol. 2, 1531–1540 (2018).

13. Smallegange, I. M., Caswell, H., Toorians, M. E. M., & de Roos, A. M. Mechanistic description of population dynamics using dynamic energy budget theory incorporated into integral projection models. Methods Ecol. Evol. 8, 146–154 (2017).

14. Augustine, S., & Kooijman, S. A. L. M. A new phase in DEB research. J Sea Res. 143, 1–7 (2019).

15. Romeijn, J., & Smallegange, I. M. Exploring how the fast-slow pace of life continuum and reproductive strategies structure microorganism life history variation. Preprint at 10.1101/2022.11.28.517963 (2022).

16. Atkinson, D., & Sibly, R. M. Why are organisms usually bigger in colder environments? Making sense of a life history puzzle. Trends Ecol. Evol. 12, 235–239 (1997).

17. Rademaker, M., van Leeuwen, A., & Smallegange, I. M. Why we should not necessarily expect life history strategies to inform on sensitivity to environmental change. Preprint at 10.22541/au.164848872.26565315/v1 (2022).

18. Kooijman, S. A. L. M., & Metz, J. A. J. On the dynamics of chemically stressed populations: The deduction of population consequences from effects on individuals. Ecotoxicol. Environ. Saf. 8, 254–274 (1984).

19. Sousa, T., Domingos, T., Poggiale, J. C., & Kooijman, S. A. L. M. Dynamic energy budget theory restores coherence in biology. Philos. Trans. R. Soc. Lond. B 365, 3413–3428 (2010).

20. Easterling, M. R., Ellner, S. P., & Dixon, P. M. Size-specific sensitivity: Applying a new structured population model. Ecology 81, 694–708 (2000).

21. Coulson, T. Integral projections models, their construction and use in posing hypotheses in ecology. Oikos 121, 1337–1350 (2012).

22. Merow, C., et al. Advancing population ecology with integral projection models: a practical guide. Methods Ecol. Evol. 5, 99–110 (2014).

23. Thunell V., Gårdmark, A., Vindenes, Y. Optimal energy allocation trade-off driven by size-dependent physiological and demographic responses to warming. Ecology 4, e3967 (2023).

24. Froese, R. Estimating somatic growth of fishes from maximum age or maturity. Acta Ichthyol. Piscat. 52, 125–133 (2022).

25. Froese, R., & Pauly, D. Editors. FishBase. World Wide Web electronic publication. https://www.fishbase.org, (02/2023).

26. Ebert, D. A., Dando, M., & Fowler, S. Sharks of the World. A complete guide (Princeton University Press, 2021).

27. Last, P., et al. Rays of the World (CSIRO Publishing, 2016).

28. IUCN. The IUCN Red List of Threatened Species. Version 2022-2. https://www.iucnredlist.org. Accessed on [19 May 2023].

29. Temple, A. J., et al. Life-history, exploitation and extinction risk of the data-poor Baraka’s whipray (Maculabatis ambigua) in small-scale tropical fisheries. J. Fish Biol. 97, 708–719 (2020).

30. Barrowclift, E., et al. Tropical rays are intrinsically more sensitive to overfishing than the temperate skates. Biol. Cons. 281, 110003 (2023).

31. Guallart, J., & Vicent, J. J. Changes in composition during embryo development of the gulper shark, Centrophorus granulosus (Elasmobranchii, Centrophoridae): an assessment of maternal-embryonic nutritional relationships. Environ. Biol. Fishes 61, 135–150 (2001).

32. Natanson, L. J., & Cailliet, G. M. Reproduction and development of the Pacific angel shark, Squatina californica, off Santa Barbara, California. Copeia 4, 987–994 (1986).

33. Simpfendorfer, C. A., & Unsworth, P. Reproductive biology of the whiskery shark, Furgaleus macki, off south-western Australia. Mar. Freshwater Res. 49, 687–793 (1998).

34. Mattos, S. M., Broadhurst, M., Hazin, F. H., & Jones, D. M. Reproductive biology of the Caribbean sharpnose shark, Rhizoprionodon porosus, from northern Brazil. Mar. Freshwater Res. 52, 745–752 (2001).

35. Pardo, S. A., Kindsvater, H. K., Reynolds, J. D., & Dulvy, N. K. Maximum intrinsic rate of population increase in sharks, rays, and chimaeras: the importance of survival to maturity. Can. J. Fish. Aquat. Sci., 73, 1159–1163 (2016).

36. Smith, S. E., Au, D. W., & Show, C. Intrinsic rebound potentials of 26 species of Pacific sharks. Mar. Freshwater Res. 49, 663–678 (1998).

37. Dulvy, N. K., et al. Methods of assessing extinction risk in marine fishes. Fish Fish. 5, 255–276 (2004).

38. Add-my-pet. Database of code, data and DEB model parameters (https://www.debtheory.org) (2023)

39. Kooijman, S. A. L. M. Dynamic energy budget theory for metabolic organization (Cambridge, UK: Cambridge University Press 2010).

40. Smallegange, I. M. DEBBIES. A database to compare life history strategies across ectotherms. Figshare. Dataset. 10.6084/m9.figshare.13241972 (2020).

41. Marques, G. M., et al. The AmP project: Comparing species on the basis of dynamic energy budget parameters. PLoS Computational Biology, 14(5), e1006100 (2018).

42. Pottor, T., Reznick, D. N., & Coulson, T. Substantial intraspecific variation in energy budgets: Biology or artifact? Funct. Ecology 35, 1693–1707 (2021).

43. Hodgson, D. J., & Townley, S. Linking management changes to population dynamic responses: the transfer function of a projection matrix perturbation. J Applied Ecol. 41, 1155–1161 (2004).

44. Deines A., et al. Robust population management under uncertainty for structured population models. Oikos 17, 2175–2183 (2007).

45. Myers, P., et al. The Animal Diversity Web (online). Accessed at https://animaldiversity.org (2023).

46. Smart, J. J., et al. Stochastic demographic analyses of the silvertip shark (Carcharhinus albimarginatus) and the common blacktip shark (Carcharhinus limbatus) from the Indo-Pacific. Fish. Res. 191, 95–107 (2017).

47. de Magalhaes, J. P., & Costa, J. A database of vertebrate longevity records and their relation to other life-history traits. J. Evol. Biol. 22, 1770–1774 (2009).

48. Caswell, H. Sensitivity Analysis: Matrix Methods in Demography and Ecology. Demographic Research Monographs. Springer Open, chapter 10 (2019).

49. Piet, G. J., & Guruge, W. A. H. P. Diel variation in feeding and vertical distribution of ten co-occurring fish species: consequences for resource partitioning. Env. Biol. Fishes 50, 293–307 (1997).

50. Revell, L. J. phytools: An R package for phylogenetic comparative biology (and other things). Methods Ecol. Evol. 3, 217–223 (2012).

51. Stearns, S. C. The influence of size and phylogeny on patterns of covariation among life-history traits in the mammals. Oikos 41, 173–187 (1983).

52. Caswell, H. Matrix Population Models. (Sunderland, MA: Sinauer Associates 2001)

53. Tuljapurkar, S., Gaillard, J. M. & Coulson, T. From stochastic environments to life histories and back. Phil. Trans R. Soc. B. 364, 1499–1509 (2009).

54. Keyfitz N., & Caswell H. Applied mathematical demography. (Springer, New York 2005).

